# Rv3737 is required for *Mycobacterium tuberculosis* growth *in vitro* and *in vivo* and correlates with bacterial load and disease severity in human tuberculosis

**DOI:** 10.1101/2021.01.04.425357

**Authors:** Qing Li, Zhangli Peng, Xuefeng Fu, Hong Wang, Zhaoliang Zhao, Yu Pang, Ling Chen

## Abstract

Rv3737 is the sole homologue of multifunctional transporter ThrE in *Mycobacterium tuberculosis* (*Mtb*). In this study, we aimed to investigate whether this transporter participates *in vitro* and *in vivo* survival of *Mtb*. To characterize the role of Rv3737, we constructed and characterized an *Mtb* H37RvΔRv3737. This strain was evaluated for altered growth rate and macrophage survival using cell model of infection. In addition, the comparative analysis was conducted to determine the association between Rv3737 mRNA expression and disease severity in active pulmonary TB patients. The H37RvΔRv3737 strain exhibited significant slow growth rate compared to H37Rv-WT strain in standard culture medium. Additionally, the survival rate of H37Rv-WT strain in macrophages was 2 folds higher than that of H37RvΔRv3737 at 72 h. A significant higher level of TNF-α and IL-6 mRNA expression was observed in macrophages infected with H37RvΔRv3737 as compared to H37Rv-WT. Of note, Rv3737 expression was significantly increased in clinical *Mtb* isolates than H37Rv-WT. The relative expression level of Rv3737 was positively correlated with lung cavity number in TB patients. Similarly, the higher Rv3737 mRNA level resulted in lower C(t) value by Xpert MTB/RIF assay, demonstrating that a positive correlation between Rv3737 expression and bacterial load in TB patients. In conclusion, our data is the first to demonstrate that the transporter Rv3737 is required for *in vitro* growth and survival of bacteria inside macrophages. In addition, the expression level of Rv3737 is associated with bacterial load and disease severity in pulmonary tuberculosis patients.

## Introduction

Tuberculosis (TB), caused by *Mtb* complex, constitutes a major global health threat. It is estimated that one third of the world’s population is latently infected by the bacterium, and 10.0 million have fallen ill with TB annually [1]. The HIV pandemic and the emergence of multidrug-resistant tuberculosis have contributed further to its spread [2, 3]. Life cycle of *Mtb* involves transition stages from infection, dormancy and reactivation, and active TB is the result of reactivation of latent TB infection in most instances [4]. Specially, individuals with immunosuppression have higher odds of TB reactivation compared with normal individuals [5]. Therefore, *Mtb* is able to survive during periods of reduced growth and has the capacity to regrow rapidly in response to stresses encountered in the host.

Bacteria are equipped with a broad variety of transport systems [6]. Transport processes play a pivotal role in pathogen metabolism, e.g. for the uptake of nutrients and excretion of harmful agents [7]. Moreover, several amino acid transporters are indeed reported to be associated with pathogenesis [6]. For instance, threonine transporter ThrE is essential for the fitness of *Corynebacterium glutamicum in vitro* growth [8]. Inactivation of *thrE* gene shows reduced growth rate *in vitro* in medium supplemented with threonine. In addition to threonine, the ThrE carrier serves to export small molecules, indicating that it is a multifunctional transporter that gets rid of metabolic waste products and thus hold more importance of *thrE* in biological fitness [8]. Of note, only two homologues are identified in *Mtb* and *S. coelicolor* [8]. Rv3737 encodes a 55 Da protein with significant sequence similarity to characterized ThrE protein [9]. As a member of new translocator family that has never been reported before, it is interesting to investigate whether this potential multifunctional transporter participates *in vitro* and *in vivo* survival of *Mtb*. To characterize the role of Rv3737, we constructed and characterized an *Mtb* H37RvΔRv3737. This strain was evaluated for altered growth rate and macrophage survival using cell model of infection. In addition, the comparative analysis was conducted to determine the association between Rv3737 mRNA expression and disease severity in active pulmonary TB patients.

## Materials and Methods

### Bacterial strains, plasmids and cells

The bacterial strains, plasmids and cells used in this study are detailed in Table S1. *E. coli* DH5α and *E. coli* HB101 cells were grown in Luria-Bertani (LB) broth or LB agar plates at 37°C. Clinical isolates of *Mtb, Mtb* reference strain H37Rv (H37Rv-WT, ATCC27294) and *Mycobacterium smegmatis* mc^2^ 155 cells growth on Lowenstein–Jensen (L-J) medium (Encode, Zhuhai, China), Middlebrook 7H9 broth or 7H10 agar plates containing 0.05% Tween 80, 0.5% glycerol and 10% OADC. The selective 7H9 broth or 7H10 agar plate supplemented with 75 μg/ml hygromycin was used to subculture *Mtb* Rv3737 knockout strain (H37RvΔRv3737). The bacteria with OD_600_ of 0.6-1.0 were used for *in vitro* experiments. RAW264.7 cultured in DMEM complete medium containing 10% Fetal Bovine Serum.

In addition, 12 clinical *Mtb* isolates were collected from a set of sputum smear-positive and GeneXpert MTB-positive specimens from August 2016 to February 2017. The demographic and clinical characteristics were obtained from electronic medical records. This study was approved by the Ethics Committees of the Affiliated Hospital of Zunyi Medical University. All the patients were ≥18 years old and provided written informed consent prior to enrolment.

### Construction of H37RvΔRv3737

Using the genomic DNA of H37Rv-WT as a template, the Rv3737 nucleic acid sequence was derived from Genbank of NCBI (https://www.ncbi.nlm.nih.gov/gene/885794). As shown in Fig. S1A, the primers for the upper and lower arms of the Rv3737 gene and verification primers were designed based on the principle of homologous recombination (Table S2). Flanking regions comprising upstream and downstream regions of the Rv3737 gene were amplified by PCR and cloned into the p0004S plasmid containing a hygromycin resistance cassette, the vector was then ligated into the phAE159 plasmid, which was electroporated into M. smegmatis, and the resulting phage was amplified to obtain a high-titer stock. The high-titer phage was used to infect H37Rv-WT, which was plated onto selective 7H10 agar plates. Plates were incubated for 4∼8 weeks at 37°C, which eventually led to the growth of a small number of H37RvΔRv3737 colonies. Colonies were picked, and PCR and qPCR was used to confirm the presence of the hygromycin-gene flanking region and the absence of the Rv3737 gene (Fig. S1B, S1C) [10, 11].

### Growth and colonial morphology of Rv3737 knockout strain

H37RvΔRv3737 was inoculated in selective 7H9 broth medium and incubated at 37°C. Optical density at 600 nm (OD600nm) was detected at intervals of 24 h and the growth curves at 37°C were obtained. H37RvΔRv3737 was inoculated in selective 7H10 agar plates. After incubation for 3 weeks at 37°C, the colony morphology was recorded with the HP scanner. The H37Rv-WT was used as a control and the same treatment was performed. For scanning electron microscope analysis, the bacteria were harvested by centrifugation at 500 rpm for 5 minutes. Then 2.5% glutaraldehyde was added into the pellet for 24 hours for fixation purpose. Followed by treatment with 1% osmic acid and ethanol gradient, the sample were sprayed with gold film and observed in a SU8010 scanning electron microscope (Hitachi, Japan)[12].

### Survival of Rv3737 knockout strain in RAW 264.7

H37RvΔRv3737 and H37Rv-WT were infected to 5 × 10^5^/well RAW 264.7 cells in 6-well plate at multiplicity of infection (MOI) 10. After 4 h of incubation, all extracellular bacteria were removed gently by washing and intracellular bacteria were harvested. At 24 h and 72h after infection, both extracellular bacteria released from macrophage lysed in supernatant and intracellular bacteria in intact cell layer were harvested. Bacteria at 4 h, 24 h and 72 h were plated on 7H10 agar plates in triplicate, plates were incubated for 3 weeks at 37°C and CFU (colony forming unit) were counted [13].

### Cytokine measurement

Culture supernatants and sediments from *Mtb*-infected RAW 264.7 cells were harvested at 0 h, 4 h, 8 h, 12 h, 24 h post-infection and stored at − 80 °C for cytokine measurement. The concentrations of TNF-α and IL-6 in culture supernatant were detected using an enzyme-linked immunosorbent assay (ELISA) kit according to the manufacturer’s instructions (Solarbio, Beijing, China)[14]. The mRNA level of TNF-α and IL-6 in Culture sediments was determined using qPCR. In simple terms, the total RNA was extracted with Trizol method according to the instructions of the manufacturers[15]. After treatment with DNaseI (TaKaRa, Dalian, China), the cDNAs were reverse-transcribed from 5 µg of total RNA with the PrimeScript™ II 1st Strand cDNA Synthesis Kit (TaKaRa, Dalian, China). Real time PCR (qPCR) was carried out in triplicates for each sample using TB Green® Premix Ex Taq™ II (TaKaRa, Dalian, China) in The CFX96 touch Real-Time PCR System (Bio-Rad) [16]. Primers for transcriptional level analysis are listed in Table S2. SigA were used as the internal control in respective qPCR experiments.

### Rv3737 expression of clinical isolates and H37Rv-WT

The *Mtb* isolates at log phage were lysed by ultrasound and then subjected to RNA extraction. The mRNA level of Rv3737 in clinical isolates and H37Rv-WT was determined using qPCR method as method above. Primers for transcriptional level analysis are listed in Table S2 and SigA were used as the internal control in respective qPCR experiments.

### Sputum collection, bacterial growth and bacterial load measurement in sputum

Sputum specimens were collected for acid-fast staining and GeneXpert MTB/RIF assay (Cepheid, Sunnyvale, CA, United States). Acid-fast staining microscopy was performed directly on all samples as described previously [17]. One milliliter sputum was mixed with 2 ml sample reagent, and incubated at room temperature for 15 min. Then the decontaminated sample was then added to a test cartridge and loaded onto the Xpert instrument. Results were reported as cycle threshold (Ct) values that represented the minimal PCR cycles required for detection threshold [18]. The average Ct values of five probes was used to estimate bacterial load after exclusion of any delayed values due to rifampicin resistance [19]. 1.0 mL of sputum specimen with positive results by both tests were treated with N-acetyl-L-cysteine-NaOH-Na citrate (2.00% final concentration). After neutralization and centrifugation, the suspension of pellet was inoculated on Lowenstein-Jensen (L-J) medium. The visible growth of colonies on L-J medium was identified using conventional biochemical method [13].

### Statistical and analysis

GraphPad Prism v7.03 (GraphPad Software, San Diego, California, USA) was used to analyze the data and generate graphs. *t* test was used to compare two groups of data, two-way ANOVA was used for three or more groups of data, considering *p* value <0.05 to be significant. The linear relationships were analyzed by the R squared correlation method. Spearman Coefficient was conducted to establish the relationship between the expression level of Rv3737 and the Ct value yielded by Xpert and between the expression level of Rv3737 and the number of cavities.

## Results

### Role of Rv3737 in the physiology of *Mtb*

At the amino acid level, Rv3737 shared 29.60% sequence identity with ThrE of *C. glutamicum* (Fig. S1). We firstly assessed whether inactivation of Rv3737 affects the *in vitro* growth and physiology of *Mtb*. As shown in Fig. 1A, the H37RvΔRv3737 strain exhibited significant slow growth rate compared to H37Rv-WT strain in standard culture medium. The colony size of H37RvΔRv3737 and H37Rv-WT strain was compared by plating the same dilution on plates after 21 days (Fig. 1B, 1C). The average colony size of H37RvΔRv3737 was 0.38±0.02 μm, which was much smaller than that of H37Rv-WT strain (0.58±0.02 μm, Fig. 1D).

**Figure 1.**
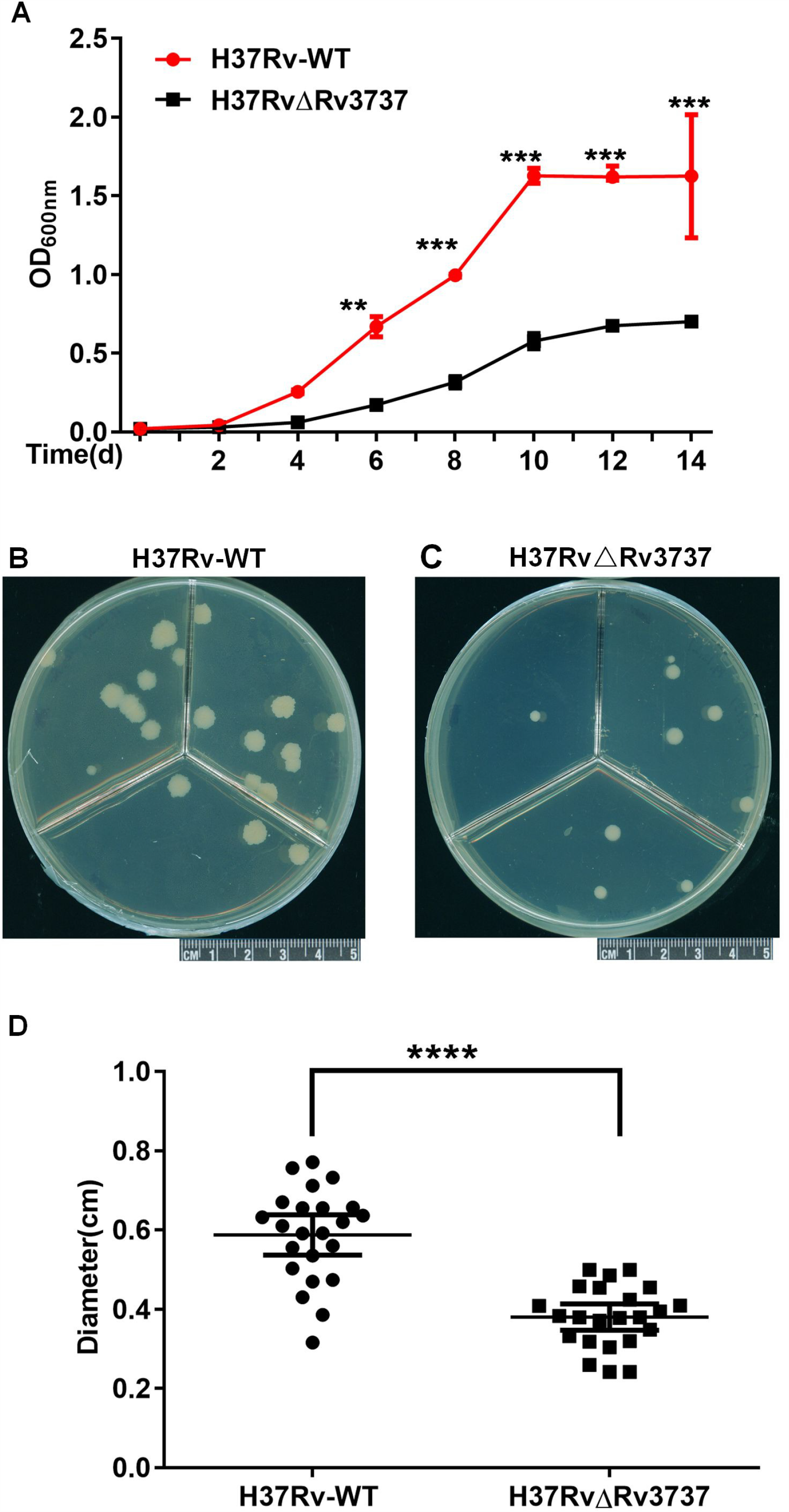
*In vitro* growth of H37RvΔRv3737. A. Growth curve of H37Rv-WT and H37RvΔRv3737 in 7H9 medium. B. Colonies of H37Rv-WT and H37RvΔRv3737 on 7H10 agar grown for 21 days. C. Diameter of colonies of H37Rv-WT and H37RvΔRv3737 on 7H10 agar. The difference between the wild type and the mutant was significant by Student’s *t* test (*, *p*<0.05; **, *p*<0.01; ***, *p*<0.001; ****, *p*<0.0001).

Field emission scanning electron microscopy was conducted to assess cell morphology and length of *Mtb*. Individuals cells from two strains were indistinguishable in morphology characteristics, whereas significant differences were observed in cell lengths. As noted in Fig. 2, the average cell length of the H37Rv-WT and H37RvΔRv3737s were 1.89 ± 0.04 and 1.69 ± 0.05 μm (*p*<0.05), respectively, suggesting that the latter had shorter cell length. In contrast, the average cell width of H37RvΔRv3737 was 0.41 ± 0.01 um, which was statistically higher than that of H37Rv-WT strain (0.35 ± 0.01, *p*<0.05).

**Figure 2.**
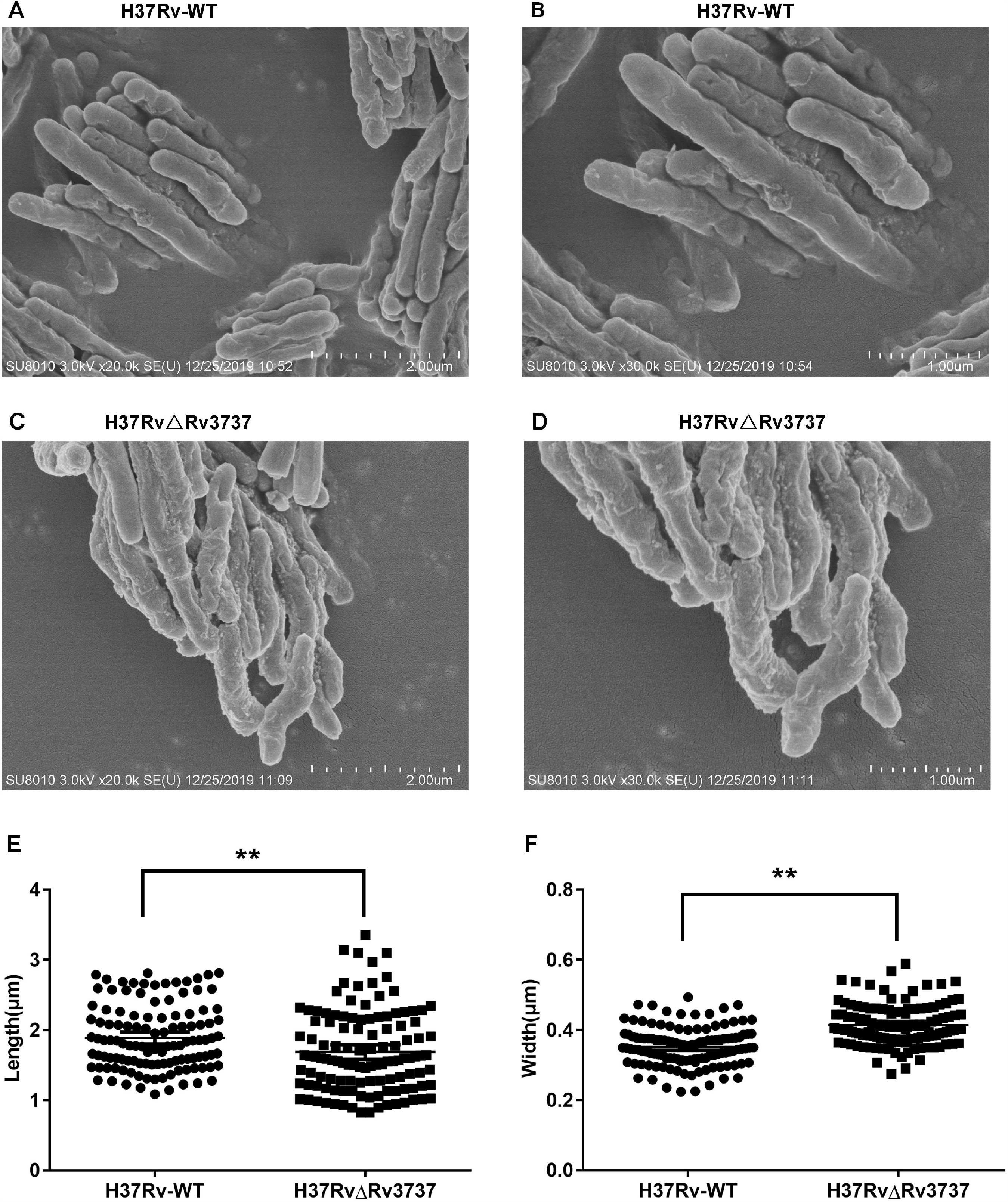
Morphological changes between the H37Rv-WT and H37RvΔRv3737 strains. A-D. Scanning electron microscope photographs of strains. Bacteria were grown in 7H9 medium. E. Length of H37Rv-WT and H37RvΔRv3737. F. Width of H37Rv-WT and H37RvΔRv3737. The difference between the wild type and the mutant was significant by Student’s t test (*, *p*<0.05; **, *p*<0.01; ***, *p*<0.001; ****, *p*<0.0001).

### Survival of H37RvΔRv3737 in macrophages

Mouse macrophages were infected with H37Rv-WT and with H37RvΔRv3737 to determine differences in capacity for intracellular growth. As shown in Fig. 3A, the infection capability of H37Rv-WT strain was approximately 2 folds higher as compared with Rv3737ΔRv3737. The intracellular survival was assessed at 24 h and 72 h after infection at 4 h as reference. The survival rate of H37Rv-WT strain was 2 folds higher than that of H37RvΔRv3737 at 72 h, respectively (Fig. 3B). Taken together, these data indicated that Rv3737 had an important role in enhancing the infection capability and intracellular survival of tubercle bacilli.

**Figure 3.**
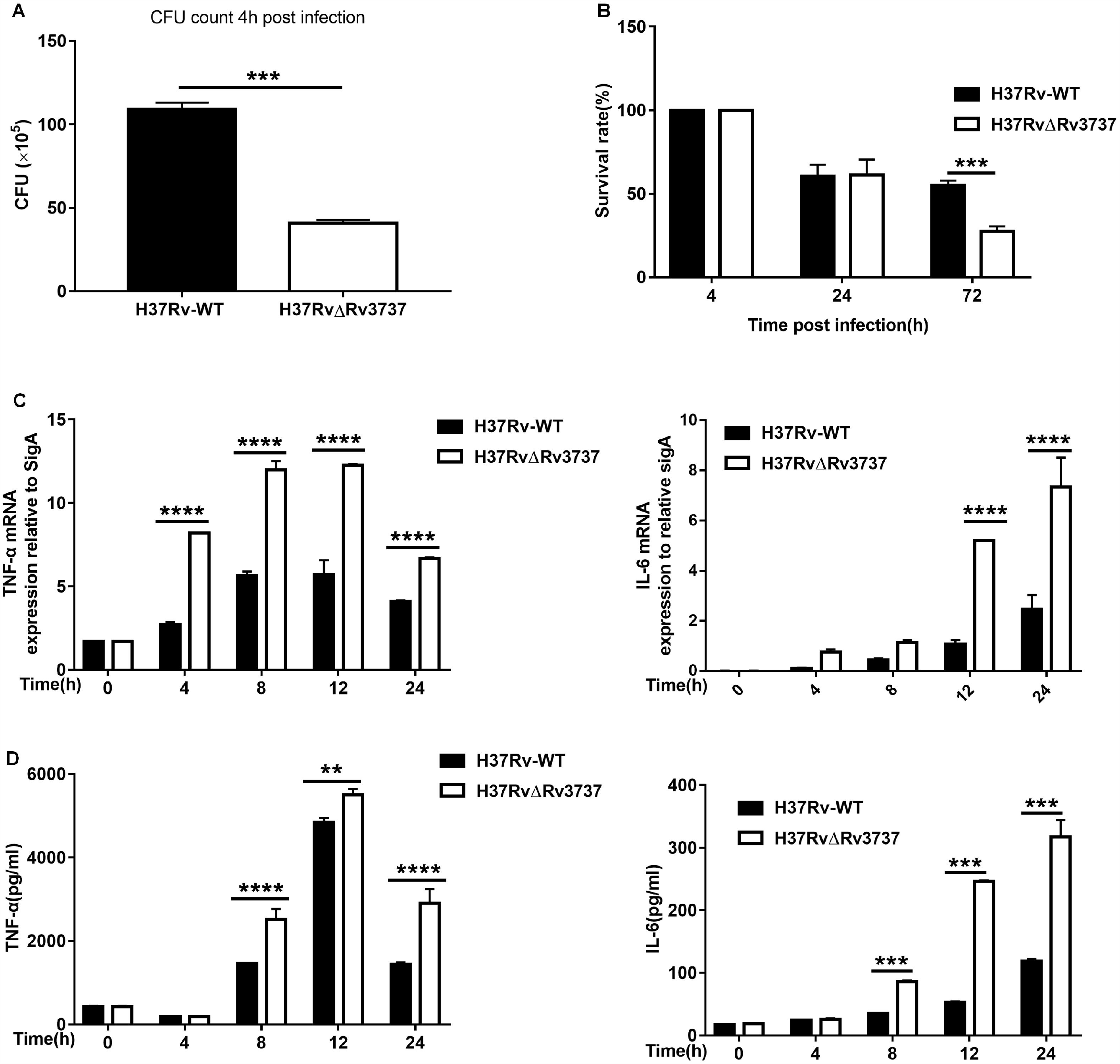
Survival of H37Rv-WT and H37RvΔRv3737 in macrophages. A. Effect of Rv3737 on infection: RAW264.7 were infected with H37Rv-WT and H37RvΔRv3737 at MOI 10 at 37 °C in CO2 incubator (5% CO2) for 4 h. B. Survival analysis of H37Rv-WT and H37RvΔRv3737 in macrophages. C. The mRNA levels of TNF-α and IL-6 in macrophages infected with H37Rv-WT and H37RvΔRv3737. D. Immunoassays for TNF-α and IL-6 in supernatants collected from macrophages infected with H37Rv-WT and H37RvΔRv3737. The difference between the wild type and the mutant was significant by Student’s *t* test (*, *p*<0.05;**, *p*<0.01; ***, *p*<0.001; ****, *p*<0.0001).

### Detection of proinflammatory cytokines in macrophages

To explore the potential role of Rv3737 in modulating the innate immune response, we investigated the levels of cytokines upon infection of RAW264.7 cells with H37Rv-WT and H37RvΔRv3737. A significant higher level of TNF-α and IL-6 mRNA expression was observed in macrophages infected with H37RvΔRv3737 as compared to H37Rv-WT (Fig. 3C). Detection of cytokines in the culture supernatants of macrophages also supported the elevated secretion of proinflammatory cytokines (TNF-α and IL-6) at increasing time points (Fig. 3D).

### Relationship between Rv3737 expression and disease severity

Based on the slower *in vivo* growth and increased host proinflammatory cytokine response, we hypothesized that the expression level of Rv3737 correlated with *Mtb* virulence in host. In order to test this hypothesis, we recruited 12 clinical *Mtb* isolates to determine whether the upregulation of Rv3737 could lead to more severe clinical symptoms. A total of 12 patients infected with diagnosed TB were retrospectively included in our analysis (Table 1). As shown in Fig. 4A, Rv3737 expression was significantly increased in clinical *Mtb* isolates than H37Rv-WT. Of note, the relative expression level of Rv3737 was positively correlated with lung cavity number in TB patients (r = 0.71, *p* < 0.01, Fig. 4B). Similarly, the higher Rv3737 mRNA level resulted in lower C(t) value by Xpert MTB/RIF assay, demonstrating that a positive correlation between R3737 expression and bacterial load in TB patients (r = −0.81, *p* < 0.01, Fig. 4C).

**Table 1.**
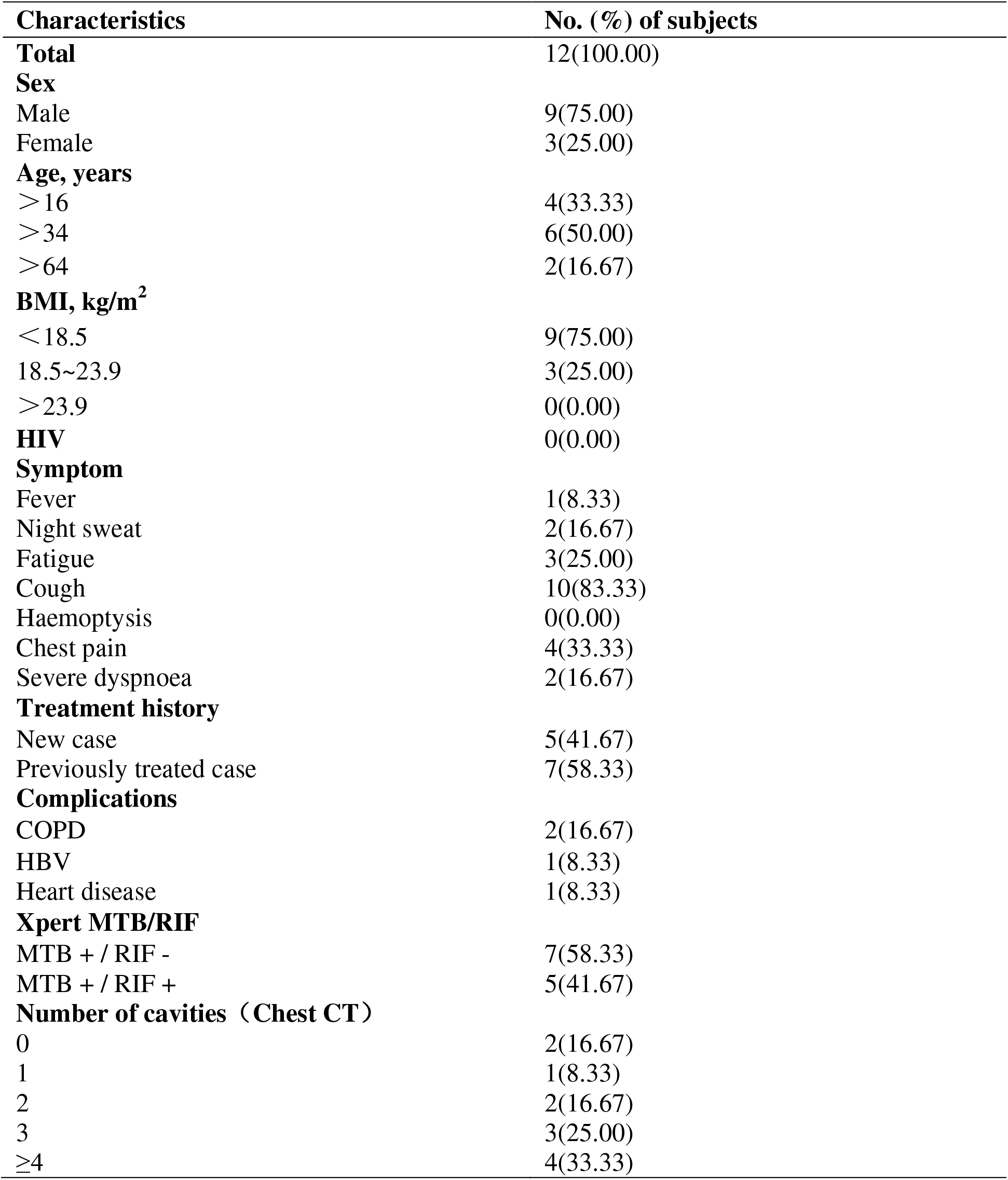
Demographic and clinical characteristics of clinical cases

| **Characteristics** | **No. (%) of subjects** |
| --- | --- |
| **Total** | 12(100.00) |
| **Sex** |  |
| Male | 9(75.00) |
| Female | 3(25.00) |
| **Age, years** |  |
| >16 | 4(33.33) |
| >34 | 6(50.00) |
| >64 | 2(16.67) |
| **BMI, kg/m<sup>2</sup>** |  |
| <18.5 | 9(75.00) |
| 18.5~23.9 | 3(25.00) |
| >23.9 | 0(0.00) |
| **HIV** | 0(0.00) |
| **Symptom** |  |
| Fever | 1(8.33) |
| Night sweat | 2(16.67) |
| Fatigue | 3(25.00) |
| Cough | 10(83.33) |
| Haemoptysis | 0(0.00) |
| Chest pain | 4(33.33) |
| Severe dyspnoea | 2(16.67) |
| **Treatment history** |  |
| New case | 5(41.67) |
| Previously treated case | 7(58.33) |
| **Complications** |  |
| COPD | 2(16.67) |
| HBV | 1(8.33) |
| Heart disease | 1(8.33) |
| **Xpert MTB/RIF** |  |
| MTB + / RIF - | 7(58.33) |
| MTB + / RIF + | 5(41.67) |
| **Number of cavities (Chest CT)** |  |
| 0 | 2(16.67) |
| 1 | 1(8.33) |
| 2 | 2(16.67) |
| 3 | 3(25.00) |
| ≥4 | 4(33.33) |

**Figure 4.**
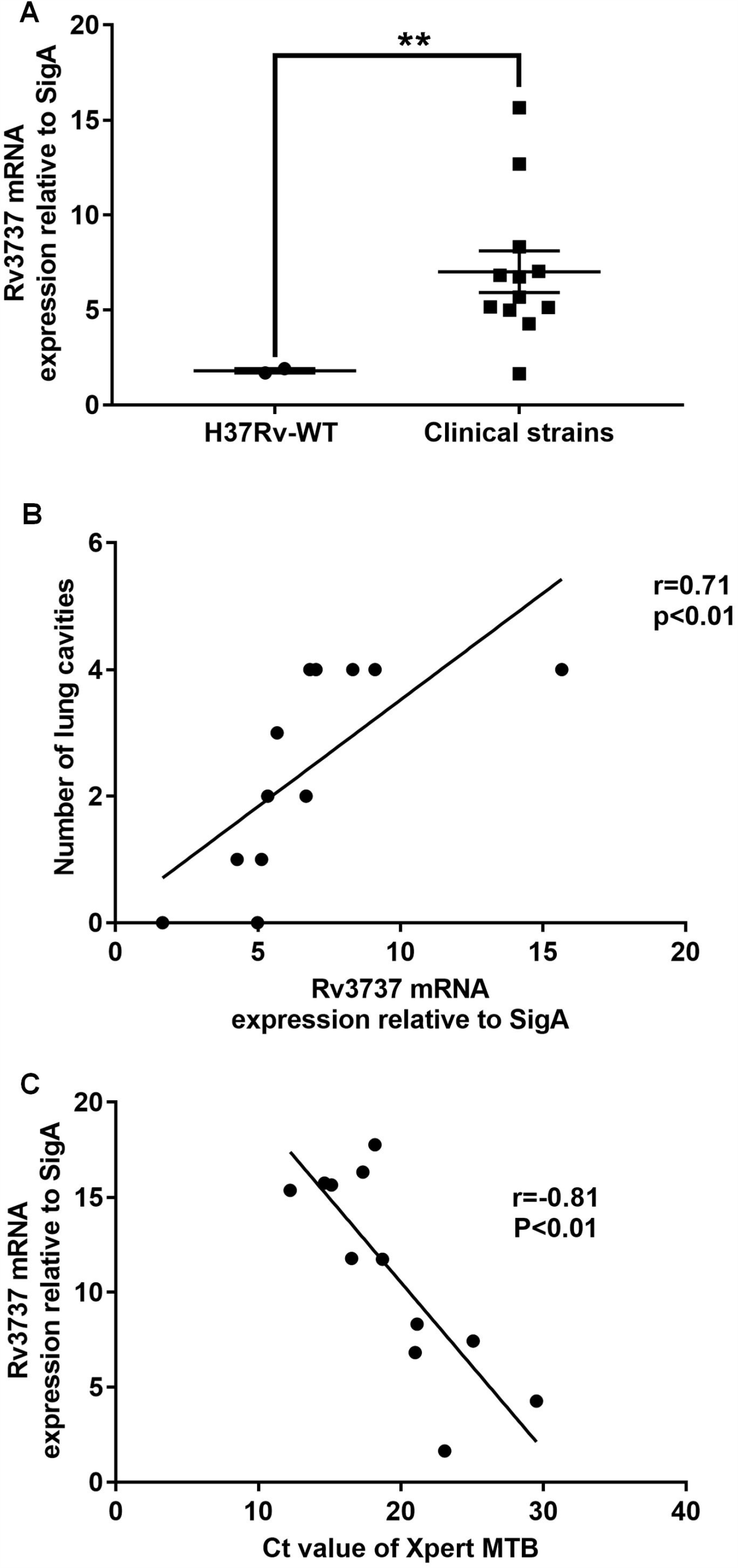
Correlation between Rv3737 mRNA level and disease severity. A. Comparison of Rv3737 mRNA levels between H37Rv-WT and clinical isolates. B. Relationship between the expression level of Rv3737 and the C(t) value yielded by Xpert. C. Relationship between the expression level of Rv3737 and the number of cavities. The difference between the wild type and the mutant was significant by Student’s t test (*, *p*<0.05; **, *p*<0.01; ***, *p*<0.001; ****, *p*<0.0001). The relationship between the expression level of Rv3737 and the C(t) value yielded by Xpert and between the expression level of Rv3737 and the number of cavities was established by Spearman Coefficient (*p* < 0.01).

## Discussion

Transporter systems are commonly considered as a potential tool for delivery of therapeutic agents. Recently experimental studies reveal that several transporters be required for chronic infection and expression of virulence in pathogenic bacteria [20, 21]. In this study, we attempted to understand the possible role of transporter protein Rv3737, which alters *in vitro* growth and intracellular survival of bacteria inside macrophages. Depletion of Rv3737 in *Mtb* resulted in decreased growth rate *in vitro* compared to H37Rv-WT; however, the mutant strain displayed usual rough and dry colonies as control strain, indicating that Rv3737 might be not involved in the cell wall lipid remodeling in *Mtb*. Although the exact characteristics and role of this transporter in growth profile remain unclear, we speculate that the inactivation of Rv3737 might lead to accumulation of metabolic waste products *in vivo* and consequently inhibit their growth. Further studies will be conducted to determine the change in metabolism profile between H37RvΔRv3737 and H37Rv-WT strain, which is essential to elucidate the substrate preference of this transporter. Experimental evidence from previous studies confirms that highly virulent *Mtb* isolates have faster *in vivo* doubling times [22, 23]. We found that the knockout of Rv3737 had a markedly negative impact on intracellular survival as compared to the control. On one hand, this fact may reflect the declined growth rate of tubercle bacilli in macrophages, as demonstrated in *in vitro* observations. On the other hand, the elevated levels of proinflammatory cytokines were noted in the culture supernatant of macrophages infected with H37RvΔRv3737, thus promoting intracellular bacteria clearance in macrophages. Specific mechanism behind this significant correlation are presently unclear. On the basis of its putative transporter function, the molecules exported by Rv3737 into extracellular substance were able to impair host defense against intracellular bacteria via inhibiting inflammatory response. On the basis of our findings, Rv3737 may participate in modulation of reduced or delayed host proinflammatory cytokine response, which is required for persisting virulence and survival of *Mtb* within host macrophages.

Furthermore, the elevated expression level of Rv3737 was noted in clinical *Mtb* isolates as compared to H37Rv-WT with attenuated virulence. This diversity supports our previous findings that Rv3737 may be involved in virulence of *Mtb*. Notably, a significant positive correlation between Rv3737 expression level and bacterial load in pulmonary TB patients raises the possibility that the isolates with increased expression of Rv3737 are prone to escaping clearance by alveolar macrophages, thus leading to greater bacillary multiplication in host. The high bacterial burden always causes more lung damage and higher mortality [24, 25]. In line with previous observation, we observed that the higher expression of Rv3737 was more likely to result in more cavities among patients affected by pulmonary TB. In view of the strong association between Rv3737 and lung pathology, we speculate that is could be used as a candidate biomarker for predicting the virulence of distinct isolates, and its potential pathogenic effect in host.

We also acknowledged several obvious limitations to the present study. First, despite exhibiting high identity in amino acid sequence with transporter ThrE, the preferential substrates of Rv3737 remains undetermined, which weaken the significance of our explanation. Second, we observed the slower growth rate and shorter length of H37RvΔRv3737; whereas the mutant had larger width than H37Rv-WT control, which may be associated with the remodeling of cytoskeleton to modulate the bacterial growth. However, the reason for this phenomenon remains unclear. Finally, although we recruited active TB cases in our analysis, the diverse courses of TB disease across patients may serve as a major challenge that biases our conclusion.

In conclusion, our data firstly demonstrate that the transporter Rv3737 is required for *in vitro* growth and survival of bacteria inside macrophages. In addition, the expression level of Rv3737 is associated with bacterial load and disease severity in pulmonary tuberculosis patients. Further studies will be conducted to determine the change in metabolism profile between H37RvΔRv3737 and H37Rv-WT strain, which is essential to elucidate the substrate preference of this transporter.

## Author contributions

QL, ZP, LC, and YP designed the experiments. QL, ZP, XF, HW, and ZZ performed the experiment.

QL, ZP, LC, and ZP wrote the manuscript. QL, ZP, CL, and YP edited and approved the manuscript.

## Acknowledgments

The study was supported by National Natural Science Foundation of China (81760003 and 81960004). The authors thank Shanghai Gene-Optimal Science & Technology Co., Ltd. for its technical support for gene knockout. The author wish to thank Prof. Mei Liu and Prof. Nana Li for collected and processed the clinical isolates. The author wish to thank Prof. Peng Xu for helpful discussions.

---

**Figure S1.**
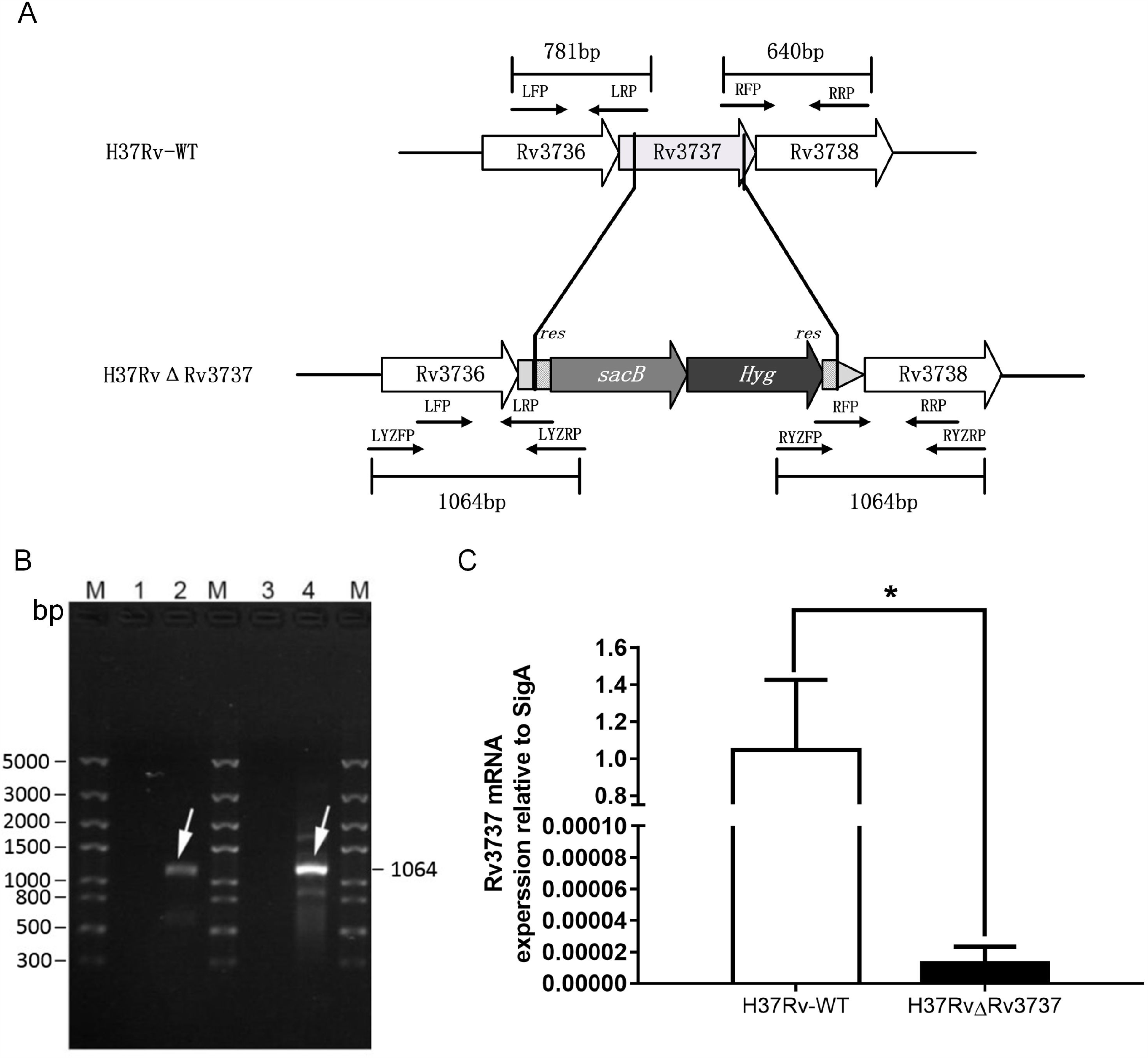
Disruption of the *Rv3737* gene in M. tuberculosis.

**Table S1.** Bacterial strains, plasmids and cells used in this study

| **Strains/plasmids/cells** | **Description** | **source/reference** |
| --- | --- | --- |
| **Strains** |  |  |
| Wild-type *Mtb* H37Rv | Pathogenic; for amplifying *M. tuberculosis* Rv3737 gene | Guizhou provincial Center For Disease Control And Prevention |
| *M.smegmatis* mc<sup>2</sup> 155 | Nonpathogenic; for amplifying *M. tuberculosis* Rv3737 gene and achieving homologous recombination at *rmlA* locus | Wuhan Institute of Virology, CAS |
| *E. coli* DH5α | For constructing plasmids | Tsingke biological technology |
| *E. coli* HB101 | For constructing plasmids | Tsingke biological technology |
| H37RvΔRv3737 | *M. tuberculosis* Rv3737 knockout strain | This work |
| *Clinical Mtb* isolates | Collected from a cohort of participants with pulmonary TB | Affiliated Hospital of Zunyi Medical University |
| **Plasmids** |  |  |
| p0004s | For constructing plasmids | Wuhan Institute of Virology, CAS |
| phAE159 | For constructing plasmids | Wuhan Institute of Virology, CAS |
| p0004s-LR | For constructing plasmids | This work |
| phAE159-p0004s-LR | For constructing plasmids | This work |
| **Cells** |  |  |
| RAW264.7 | Mouse mononuclear macrophage leukemia cells | Affiliated Hospital of Zunyi Medical University |

**Table S2.**
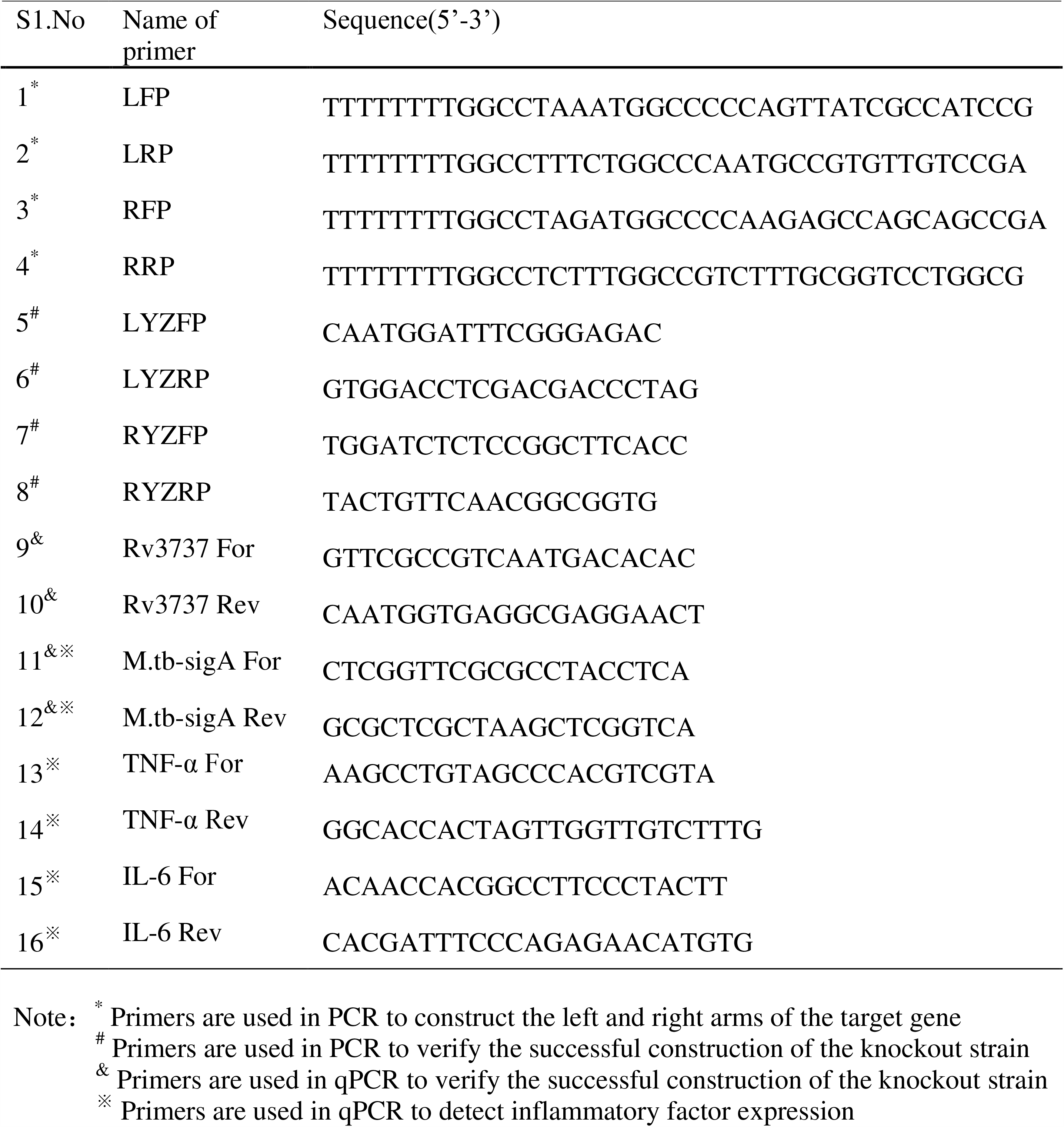
List of primer used in this study

